# Can we obtain *in vivo* transmural mean hoop stress of the aortic wall without knowing patient-specific material properties and residual deformations?

**DOI:** 10.1101/366849

**Authors:** Minliang Liu, Liang Liang, Haofei Liu, Ming Zhang, Caitlin Martin, Wei Sun

## Abstract

It is well known that residual deformations/stresses alter the mechanical behavior of arteries, e.g. the pressure-diameter curves. In an effort to enable personalized analysis of the aortic wall stress, approaches have been developed to incorporate experimentally-derived residual deformations into *in vivo* loaded geometries in finite element simulations using thick-walled models. Solid elements are typically used to account for “bending-like” residual deformations. Yet, the difficulty in obtaining patient-specific residual deformations and material properties has become one of the biggest challenges of these thick-walled models. In thin-walled models, fortunately, static determinacy offers an appealing prospect that allows for the calculation of the thin-walled membrane stress without patient-specific material properties. The membrane stress can be computed using forward analysis by enforcing an extremely stiff material property as penalty treatment, which is referred to as the forward penalty approach. However, thin-walled membrane elements, which have zero bending stiffness, are incompatible with the residual deformations, and therefore, it is often stated as a limitation of thin-walled models. In this paper, by comparing the predicted stresses from thin-walled models and thick-walled models, we demonstrate that the transmural mean hoop stress is the same for the two models and can be readily obtained from *in vivo* clinical images without knowing the patient-specific material properties and residual deformations. Computation of patient-specific mean hoop stress can be greatly simplified by using membrane model and the forward penalty approach, which may be clinically valuable.

## 1 Introduction

Residual deformations/stresses first discovered in the 1980s (Chuong and Fung 1986; Vaishnav and Vossoughi 1983) have been shown to significantly affect the physiological wall stress distributions (Delfino et al. 1997; Fung 1991; Holzapfel et al. 2000; Humphrey 2002; Matsumoto and Hayashi 1996). To incorporate residual deformations in arteries, traditional forward analysis uses a thick-walled model starting from the stress-free reference configuration. Then deformation relations, constitutive laws and equilibrium equations are utilized to solve the boundary value problem. However, when applying this conventional approach to obtain patient-specific stress fields from the *in vivo* loaded geometries in clinical images, one has to first determine the unknown material parameters and residual deformations, which are required in the thick-walled finite element (FE) models. Some studies have suggested the use of experimentally-determined material and residual deformation parameters (Alastrué et al. 2010; Pierce et al. 2015). However, using residual deformations and material properties that are not patient-specific is a clear limitation.

Fortunately, for a specific type of biological membrane structures such as the aorta, the wall stress is nearly insensitive to the variation of material properties. This property is called static determinacy, i.e. the external force (pressure) along the geometry can be used to directly compute the internal tension/stress. This is because the vessel wall can be seen as locally in a plane stress state (Miller and Lu 2013), the solution of the equilibrium is weakly sensitive to the material properties. The aorta is shown to be approximately statically determinate (Joldes et al. 2016; Liu et al. 2017). Thus, its stress distribution can be directly obtained using membrane elements by a forward penalty method (Joldes et al. 2016; Lu and Luo 2016) which enforces an extremely stiff material property as penalty treatment. The computation of the thin-walled stress can be greatly simplified by this forward approach. However, due to the assumption of no bending stiffness in the membrane elements, the self-equilibrium residual deformations are inadmissible to the thin-walled models, which is often stated as a limitation of such models.

In this paper, by comparing the predicted stresses from thin-walled models with thick-walled models considering residual deformations, we demonstrate that the transmural mean hoop stress (i.e., averaged stress through the thickness) fields are the same for the two models. Thus, the transmural mean hoop stress can be readily obtained from *in vivo* clinical images using the forward penalty approach without knowing the patient-specific material properties and residual deformations. The remaining sections are organized as follows. In Section 2, the theoretical arguments are described and the validity is shown by analytical examples. Thin-walled and thick-walled FE models with a patient-specific geometry are demonstrated in Section 3. In Section 4, the discussion and conclusions are presented.

## 2 Theoretical and Analytical Arguments

One prominent example of static determinacy is the use of Laplace law to compute the wall hoop stress by assuming a perfect cylindrical shape of the aorta.

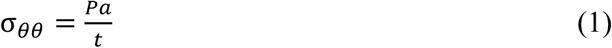

where *σ_θθ_* is the hoop stress in the thin-walled tube, *P* is the pressure, *a* is the inner radius, and *t* is the *in vivo* wall thickness. The material properties are not involved in this equation, and stress is directly calculated using the static force equilibrium. Opposite to the middle radius value used in (Horný et al. 2014), we emphasize that inner radius should be used as the blood pressure is applied to the inner surface of the aorta.

It is well known that residual stresses alter the mechanical response of arteries, e.g. the pressure-diameter curves (Holzapfel et al. 2000). Nonetheless, from the static determinacy prospective, for the *in vivo* loaded configuration, the equilibrium between the resultant force and the external pressure load should always hold, and thus, the hoop stress resultant (tension) should be insensitive to the material parameters and residual deformations. This implies that no matter how the aorta is internally balanced or residually stressed, the wall tension can always be computed only using the static equilibrium. Therefore, when the wall thickness is given, the simple thin-walled model would be sufficient in determining the transmural mean hoop stress.

### 2.1 The Opening Angle Method

In this subsection, we use an analytical example to demonstrate that the mean hoop stress is insensitive to the change of opening angles. We assume that the residual stress can be described by the opening angle and that the aorta can be modelled as a perfect tube.

Starting from the cut-open, stress-free configuration, summarizing from (Holzapfel et al. 2000) and (Pierce et al. 2015), the total deformation gradient tensor of the tube taking into account the residual stress, ***F****_total_*, can be obtained as

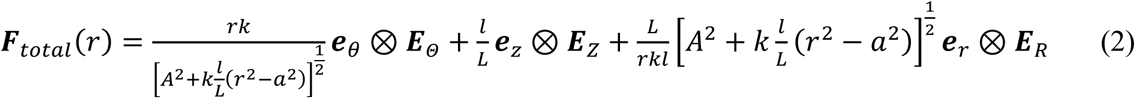

where *r ∈ [a, b*], *a* and *b* are the inner and outer radii of the *in vivo* deformed geometry. *k*, defined as 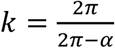, is used to describe the opening angle *α. A* and *B* are the inner and outer radii of the stress-free geometry. *L* and *I* are the axial length of the aorta segment in the stress-free and deformed geometry, respectively. ***E****_Θ_*, ***E****_Z_* and ***E****_R_* and ***e****_θ_*, ***e****_z_* and ***e****_r_* are the unit basis vectors for the stress-free and deformed geometry respectively. To make the solution simple, the constitutive relation of the aorta tissue is modelled using the isotropic Neo-Hookean model. The strain energy Ψ is

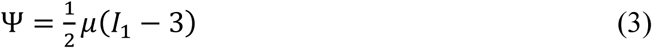

where *μ* is the shear modulus and *I*_1_ is the first invariant. To solve for the *in vivo* stress when systolic blood pressure *(P* = 104*mmHg)* (Martin et al. 2015) is present, we utilize the stress equilibrium equation, which can be expressed in the radial equation

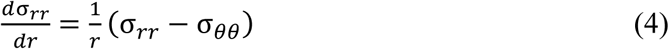

where *σ_θθ_* and *σ_rr_* are the stresses in the circumferential and radial direction respectively. Eqn. (4) can be reduced to 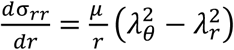 (Holzapfel and Ogden 2010), with *λ_θ_* and *λ_r_* referring to the stretches in the circumferential and radial directions, respectively. By solving the equilibrium Eqn. (4), together with the traction continuity condition *σ_rr_ (a) = −P,* we are able to obtain the radial stress (Holzapfel and Ogden 2010)

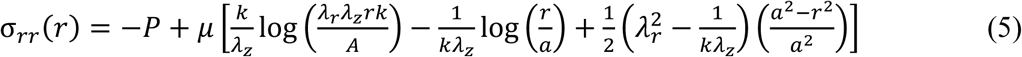

where *λ_z_* is the stretch in the axial direction. The hoop stress is then calculated using

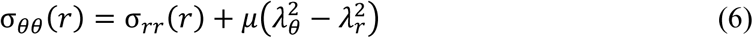

The geometry of the aorta in clinical images is always in the *in vivo* deformed state, from which the opening angle is not measurable. To this end, we fixed the inner and outer radii of the *in vivo* deformed geometry, *a* and *b,* for all scenarios and vary the opening angle from 0 to 330 degree. For a certain opening angle *α,* the inner and outer radii *A* and *B* of the cut-open sectors are solved using the boundary condition *α_rr_ (b*) = 0 and the assumption of incompressibility 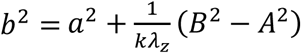. Related parameters are listed in Table 1 and values of *A* and *B* are shown in Table 2. The transmural mean hoop stress is defined as

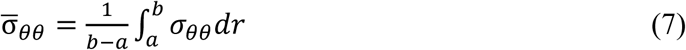

**Table 1.**
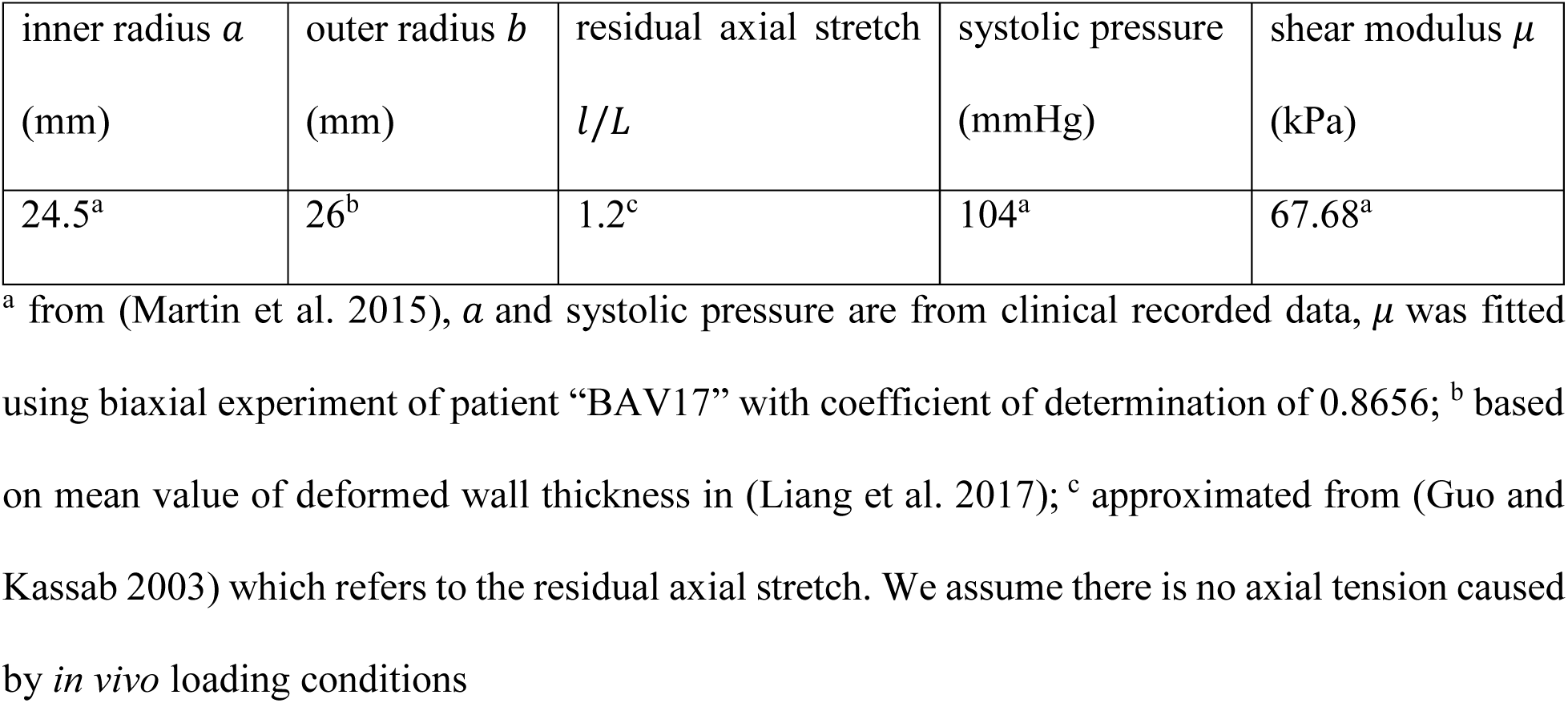
the parameters used in the opening angle method.

| inner radius $a$<br>(mm) | outer radius $b$<br>(mm) | residual axial stretch<br>$l/L$ | systolic pressure<br>(mmHg) | shear modulus $\mu$<br>(kPa) |
| --- | --- | --- | --- | --- |
| 24.5<sup>a</sup> | 26<sup>b</sup> | 1.2<sup>c</sup> | 104<sup>a</sup> | 67.68<sup>a</sup> |

<sup>a</sup> from (Martin et al. 2015), $a$ and systolic pressure are from clinical recorded data, $\mu$ was fitted using biaxial experiment of patient “BAV17” with coefficient of determination of 0.8656; <sup>b</sup> based on mean value of deformed wall thickness in (Liang et al. 2017); <sup>c</sup> approximated from (Guo and Kassab 2003) which refers to the residual axial stretch. We assume there is no axial tension caused by *in vivo* loading conditions

The results are shown in Figure 1 (left). The mean hoop stress computed from Eqn. (7) is exactly the same as the thin-walled hoop stress calculated using Eqn. (1). Unsurprisingly, if we rewrite Eqn. (4) as 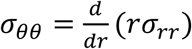 and therefore 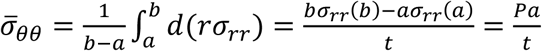, which is exactly the same formula as the Laplace law. In addition, as shown in Figure 1 (right), the adoption of anisotropic constitutive model (the GOH model (Gasser et al. 2006), described in Section 3.2.1, parameters shown in Table 3) would not affect the static determinacy. The inner hoop stress tends to be reduced while the outer hoop stress is increased when gradually increasing the opening angle. Here, small opening angles may be unusual to observe in experiment (Sokolis 2015), they are presented here for illustrative purpose.

**Table 2.** the inner and outer radii *A* and *B* of the stress-free configurations corresponding to various opening angles *α.*

| Neo-Hookean Model |  |  |  |  |  |  |  |
| --- | --- | --- | --- | --- | --- | --- | --- |
| $\alpha(^{\circ})$ | 0 | 60 | 90 | 120 | 180 | 270 | 330 |
| $A(mm)$ | 11.55 | 14.20 | 15.96 | 18.16 | 24.77 | 51.23 | 157.10 |
| $B(mm)$ | 14.98 | 17.62 | 19.39 | 21.59 | 28.20 | 54.67 | 160.54 |
| Gasser-Ogden-Holzapfel (GOH) Model |  |  |  |  |  |  |  |
| $\alpha(^{\circ})$ | 0 | 60 | 90 | 120 | 180 | 270 | 330 |
| $A(mm)$ | 18.25 | 22.12 | 24.70 | 27.93 | 37.63 | 76.43 | 231.66 |
| $B(mm)$ | 20.59 | 24.46 | 27.04 | 30.26 | 39.97 | 78.77 | 234.00 |

**Table 3.** GOH material parameters of the patient “BAV17” extracted from (Martin et al. 2015). Coefficient of determination of the curve fitting is 0.9551.

| $C_{10}(kPa)$ | $k_1(kPa)$ | $k_2$ | $\kappa$ | $\theta(^{\circ})$ |
| --- | --- | --- | --- | --- |
| 27.91 | 512.56 | 0.00 | 0.31 | 90.00 |

**Figure 1.**
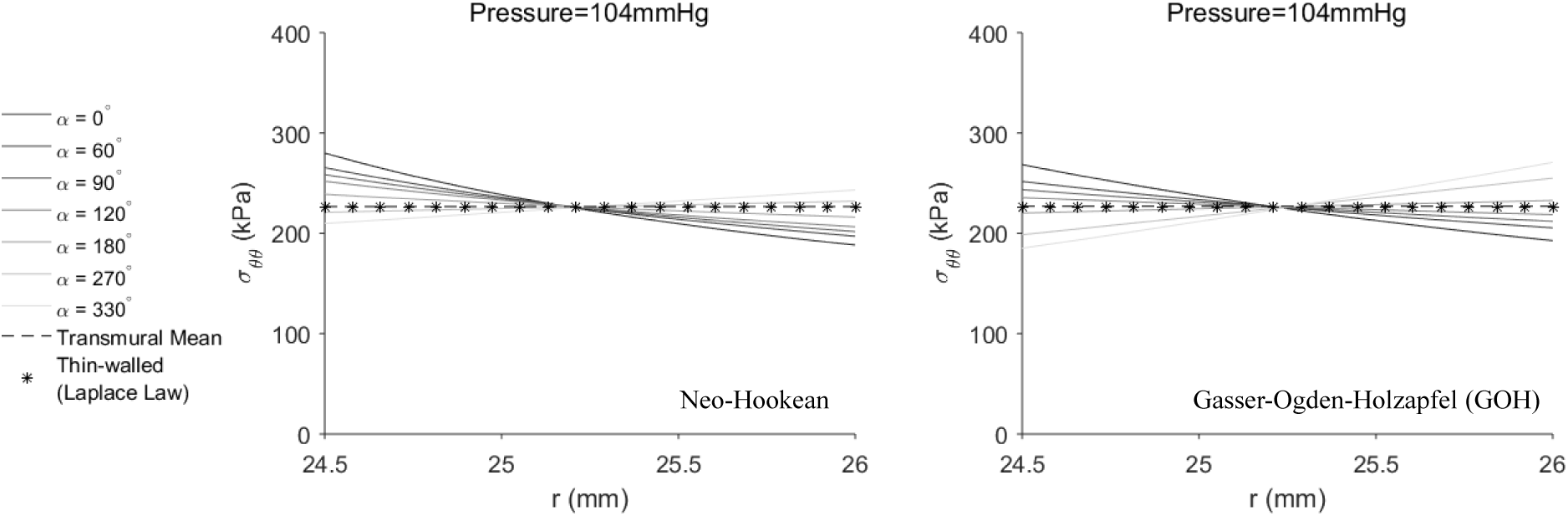
the transmural mean, thin-walled and thick-walled hoop stresses across the wall thickness. In the left figure, thick-walled hoop stresses were computed using Neo-Hookean model, while in the right figure, GOH model was used. Transmural mean hoop stress remains the same for all scenarios, thus only one line is plotted.

### 2.2 The Layer-Specific Three-Dimensional Residual Stress Model

As a step forward, (Holzapfel and Ogden 2010) proposed a layer-specific three-dimensional residual stress model, in which the residual deformations (stretching and bending) of the three layers (intima, media and adventitia) from (Holzapfel et al. 2007) were encompassed and the residual stresses were calculated using the isotropic Neo-Hookean model. In this section, we first replicate the stress distribution in (Holzapfel and Ogden 2010) by using the original parameters of geometry, material and residual deformations. Next, physiological pressure is applied to the residually-stressed aorta. The result indicates that the transmural mean hoop stress is independent of residual deformations.

The deformation gradient tensors for intima (I), media (M) and adventitia (A) (Pierce et al. 2015) are

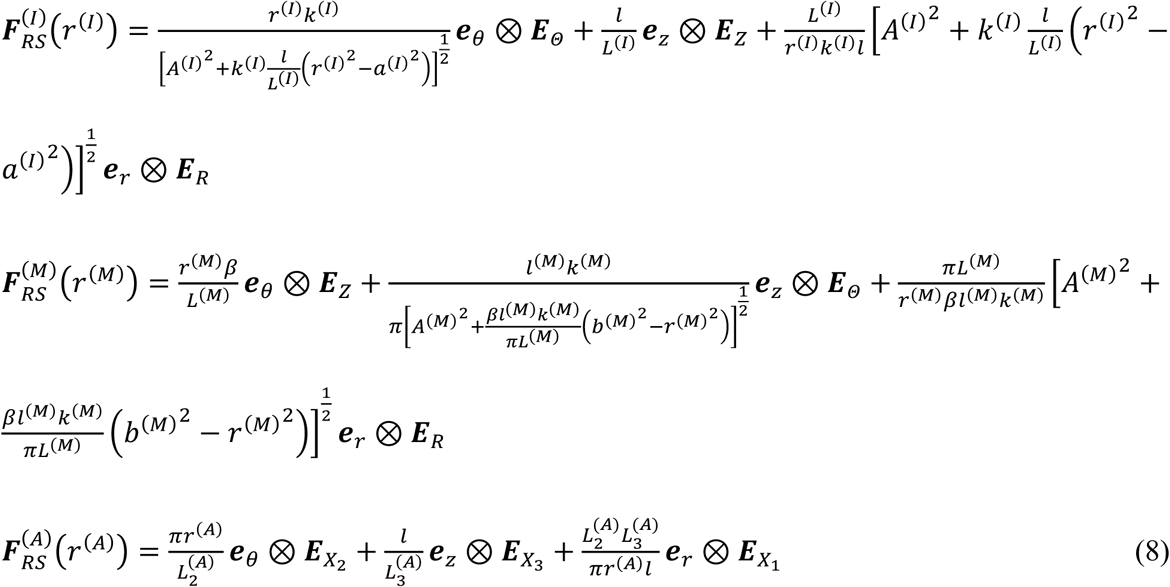

The definitions and values of the parameters are referred to (Holzapfel and Ogden 2010). Values of the related parameters are listed in Table 4.

**Table 4.** material and residual deformation parameters from (Holzapfel and Ogden 2010; Holzapfel et al. 2007). In addition, *l = 2.48mm, b^(I)^ = a^(M)^* and *b^(M)^ = a^(A)^* can be calculated according to (Holzapfel and Ogden 2010).

| Intima | Media | Adventitia |
| --- | --- | --- |
| $A^{(I)} = 7.50mm$<br>$B^{(I)} = 7.76mm$<br>$L^{(I)} = 2.58mm$<br>$k^{(I)} = 1.19$<br>$a^{(I)} = 5.61mm$<br>$\mu^{(I)} = 39.8kPa$ | $A^{(M)} = 8.41mm$<br>$B^{(M)} = 8.99mm$<br>$L^{(M)} = 2.52mm$<br>$k^{(M)} = 2.79$<br>$\mu^{(M)} = 31.4kPa$ | $L_1^{(A)} = 0.21mm$<br>$L_2^{(A)} = 18.35mm$<br>$L_3^{(A)} = 2.29mm$<br>$b^{(A)} = 7.05mm$<br>$\mu^{(A)} = 17.3kPa$ |

Similar to the procedures for the opening angle method, the hoop stress can be computed using the equilibrium equation and the boundary conditions. Interested readers are referred to (Holzapfel and Ogden 2010) for details. A diastolic pressure (*P* = 80*mmHg)* is applied to the inner surface of the aorta, and we assume no axial tension caused by *in vivo* loading conditions. The residual axial stretches have been incorporated in the deformation gradient tensors of each layer. The transmural mean hoop stress for the three layer composite is defined as

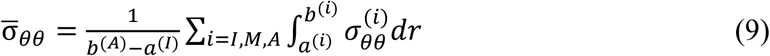

As depicted in Figure 2, the mean hoop stress is identical to the thin-walled hoop stress.

**Figure 2.**
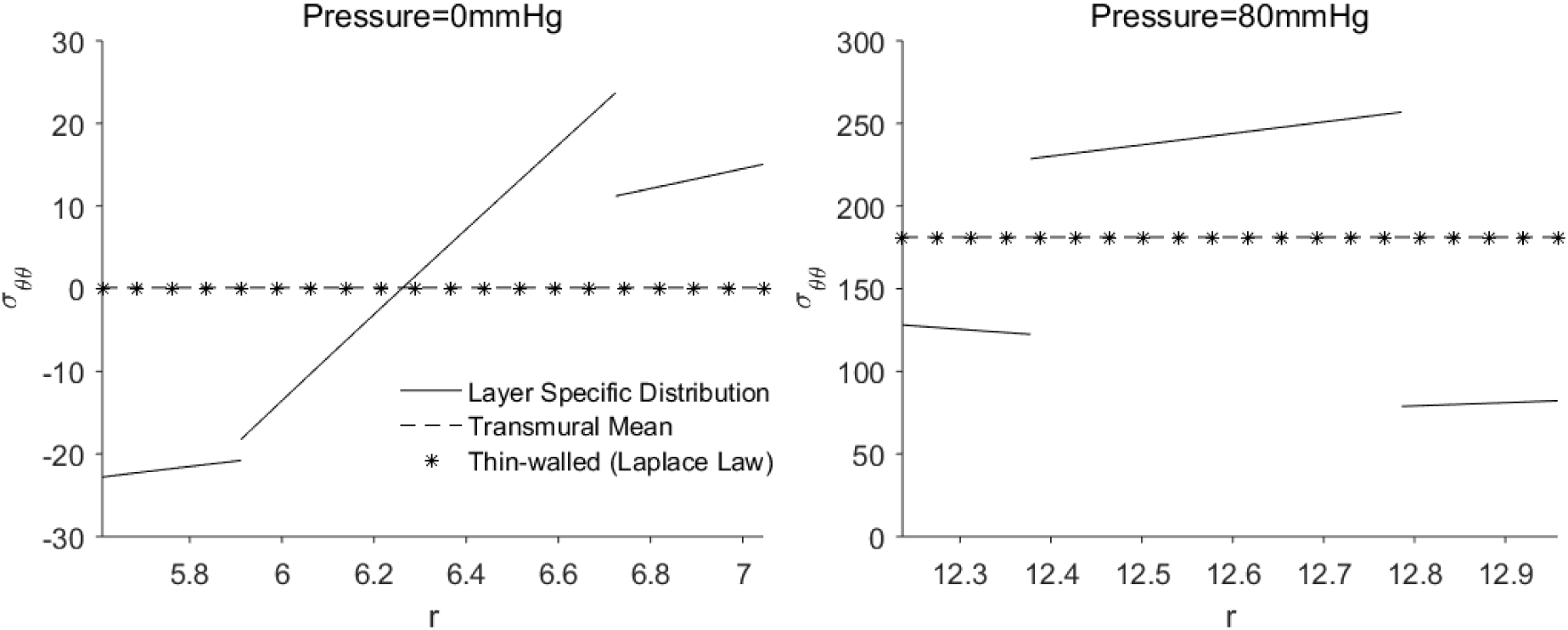
the transmural mean, thin-walled and layer-specific hoop stress distributions in the three layer composite wall when 0 and 80 mmHg pressures are applied.

## 3 Finite Element Analyses Incorporating Residual Deformations

In this section, irregularity of patient-specific geometries are taken into account using FE analyses. The validity of the conclusion in Section 2 is examined by a real patient geometry. The forward penalty approach (Section 3.1) is used to estimate the thin-walled membrane stress. For the thick-walled FE models, the generalized pre-stressing algorithm (GPA) (Pierce et al. 2015; Weisbecker et al. 2014) is implemented in ABAQUS (Section 3.2) to predict the *in vivo* stress distribution with both the residual deformations and the pre-stresses incorporated.

A CT-derived geometry from the ascending thoracic aortic aneurysm (ATAA) patient (Martin et al. 2015) were used. The inner surface of the aortic wall was divided into 4,950 M3D4 membrane elements in ABAQUS, using the automatic algorithm (Liang et al. 2017) previously developed by our group. Mesh sensitivity analysis was performed in our previous work (Martin et al. 2015). Due to partial volume effect, the wall thickness is difficult to infer from CT images, therefore a constant deformed thickness of 1.5 mm was assumed based on (Liang et al. 2017). Sensitivity analyses with respect to the wall thickness were carried out in Section 3.2.2. Next, the membrane mesh was extruded outwardly to create two solid meshes (C3D8 elements) with 8 and 9 layers in Section 3.2.2 and Section 3.2.3, respectively.

### 3.1 A Thin-walled Model using the Forward Penalty Approach

The prediction of the *in vivo* stress of the aortic wall has been relied on the recovery of the unloaded state and the incorporation of residual deformations, which requires the use of iterative techniques (Alastrué et al. 2010). A simple and effective forward penalty approach (Joldes et al. 2016) (Lu and Luo 2016) has been recently proposed to predict the *in vivo* membrane stress without knowing the material properties. In statically determinant structures, the stress is independent of the material properties, it would be legitimate to assume an extremely stiff property, so that the deformation/change of shape from the unloaded configuration to the loaded configuration is infinitesimal/negligible. This allows us to use the *in vivo* configuration as the unloaded configuration because the deformation is infinitesimal. In the forward method, an artificially stiff material property (i.e. *μ* = 10^7^*Pa,* Neo-Hookean model) is assigned to the aortic wall, realizing a penalty treatment to enforce a nearly rigid condition (Lu and Luo 2016). When the *in vivo* pressure is applied to the *in vivo*, image-derived geometry, the deformation would be infinitesimal due to the high stiffness of the material. The correct *in vivo* membrane stress field is readily obtained in this forward analysis due to the fact that the aortic wall is approximately statically determinate. This approach was shown as effective as iterative approach (Lu and Luo 2016).

Similar to the reason for the use of the inner radius in the Laplace equation (Eqn. (1)), we emphasize that the inner surface of the aortic wall should be used in the thin-walled model when applying the forward approach.

### 3.2 Thick-walled Models Incorporating Residual Deformations

#### 3.2.1 Method to incorporate Residual Deformations to Thick-walled Models

The aortic tissue is described by the Gasser-Ogden-Holzapfel (GOH) model (Gasser et al. 2006)

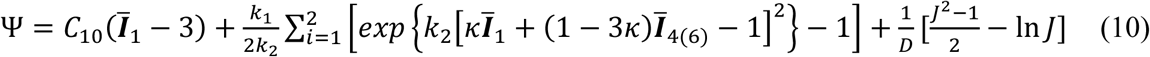

where *C*_10_, *k*_1_, *k*_2_ and *κ* are material parameters, *θ* defines the fiber directions, please refer to (Abaqus 2014; Gasser et al. 2006) for detailed definitions. The parameter *D* enforces the nearly incompressibility and is fixed to be 1 × 10^−5^.

The GPA (Pierce et al. 2015; Weisbecker et al. 2014) is utilized to incorporate the residual deformation. The total deformation gradient ***F****_t_* is stored as a history variable for each integration point. ***F****_t_* is updated based on the incremental deformation gradient *Δ****F*** resulting from the prescribed load and boundary conditions.

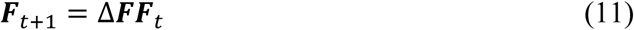

The incremental deformation gradient of the residual stress *Δ****F****_RS_* is first iteratively applied to the image-derived geometry and stored in ***F****_t_*. Next, the incremental deformation gradient of the pre-stress *Δ****F****_PS_* resulting from the *in vivo* blood pressure is incrementally applied and stored in ***F****_t_*. Thus, deformation gradient tensors associated with the residual stress ***F****_RS_* and the pre-stress ***F****_PS_* are accounted sequentially. The GPA is implemented in the ABAQUS user subroutine UMAT. The implementation was validated by comparing the analytical and FE results as in (Pierce et al. 2015).

#### 3.2.2 Thick-walled Models with Various Opening Angles

The thick-walled solid elements were utilized in this section to encompass the opening angle. Various values of the opening angle were incorporated through the GPA. Small opening angles may be unusual to observe in experiments, they are shown here for illustration purposes. The aorta was modelled as a single layer wall. This assumption may be relevant to abdominal aneurysmal tissue since collagen structure becomes nearly homogenous across the entire wall (Gasser et al. 2012). For ascending aortic aneurysms, collagen organization may be different in different layers (Sassani et al. 2015). The GOH model (Eqn (10)) was used as the constitutive law, and the material parameters (shown in Table 3) were determined from fitting the biaxial data from (Martin et al. 2015) of the particular patient.

Mean absolute percentage error (MAPE) was used to compare the transmural mean hoop stress (Eqn (7)) of the thin-walled and thick-walled models:

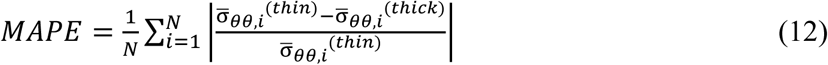

where 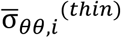 and 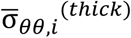 are the transmural mean hoop stress predicted by the thin-walled and thick-walled models respectively. *i* is an element index for the thin-walled model and *N* is the number of elements.

To study the sensitivity of the MAPE of the transmural mean stress with respect to the thickness, three representative thickness values (1mm, 2mm and 3mm) were chosen with *α =* 120°. The results are summarized in Table 5. Note that this opening angle value is chosen because the corresponding stress distribution is close to homogenized state in the FE simulation, and this value may not be consistent with the average value obtained from experiment (Sokolis 2015). We also notice that opening angle values are widely distributed according to (Sokolis 2015), 120 degree can be considered as a feasible value.

**Table 5.** sensitivity of MAPE w.r.t. the thickness.

| Wall thickness (mm) | 1.0 | 2.0 | 3.0 |
| --- | --- | --- | --- |
| MAPE | 0.0191 | 0.0355 | 0.0458 |

In order to quantify the transmural variation, we define a signed transmural percentage error (STPE), corresponding to the *i*th thin-walled membrane element, as

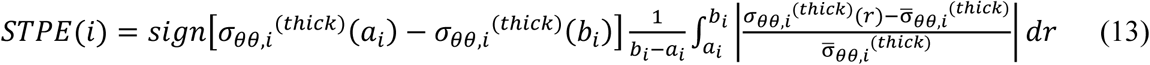

where *a_i_* and *b_i_* represent the inner and outer radii respectively, and *r* is the radius. The sign is given based on the difference between the inner and outer wall hoop stress. If the inner wall stress is greater than the outer, the STPE is positive, otherwise the STPE is negative.

The results are shown in Figure 3, the transmural mean hoop stress fields are almost identical for various opening angles and the forward penalty approach. More detailed views of ring cuts at the same location are shown in Figure 4. With increased opening angle, the mean signed transmural percentage error (MSTPE) changes from positive to negative.

**Figure 3.**
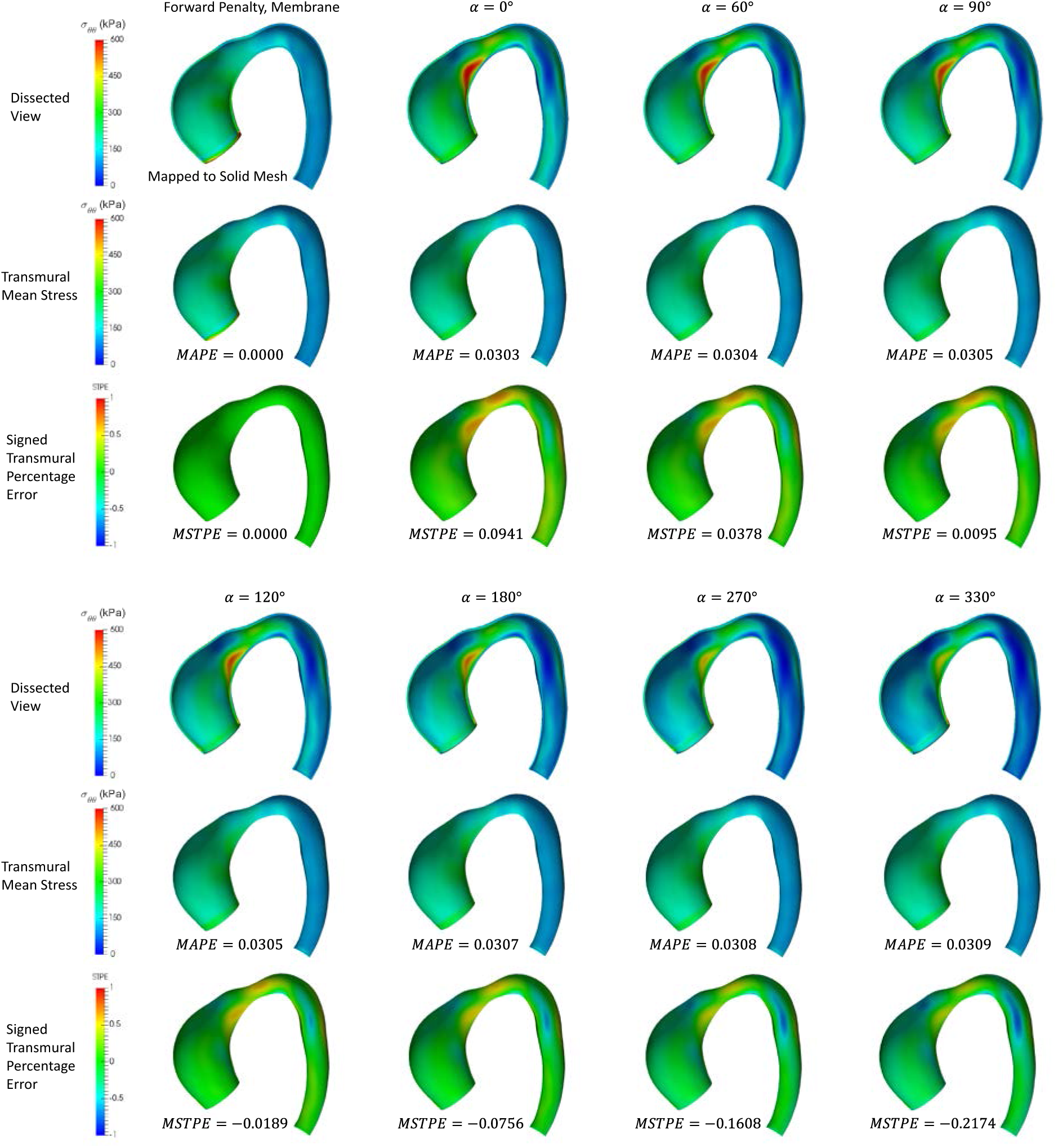
predicted results using the forward penalty approach and the GPA approach with different opening angles: (1) the hoop stress distribution in the dissected view (row 1 and row 4), (2) the transmural mean hoop stress (row 2 and row 5), and (3) the signed transmural percentage error (row 3 and row 6).

**Figure 4.**
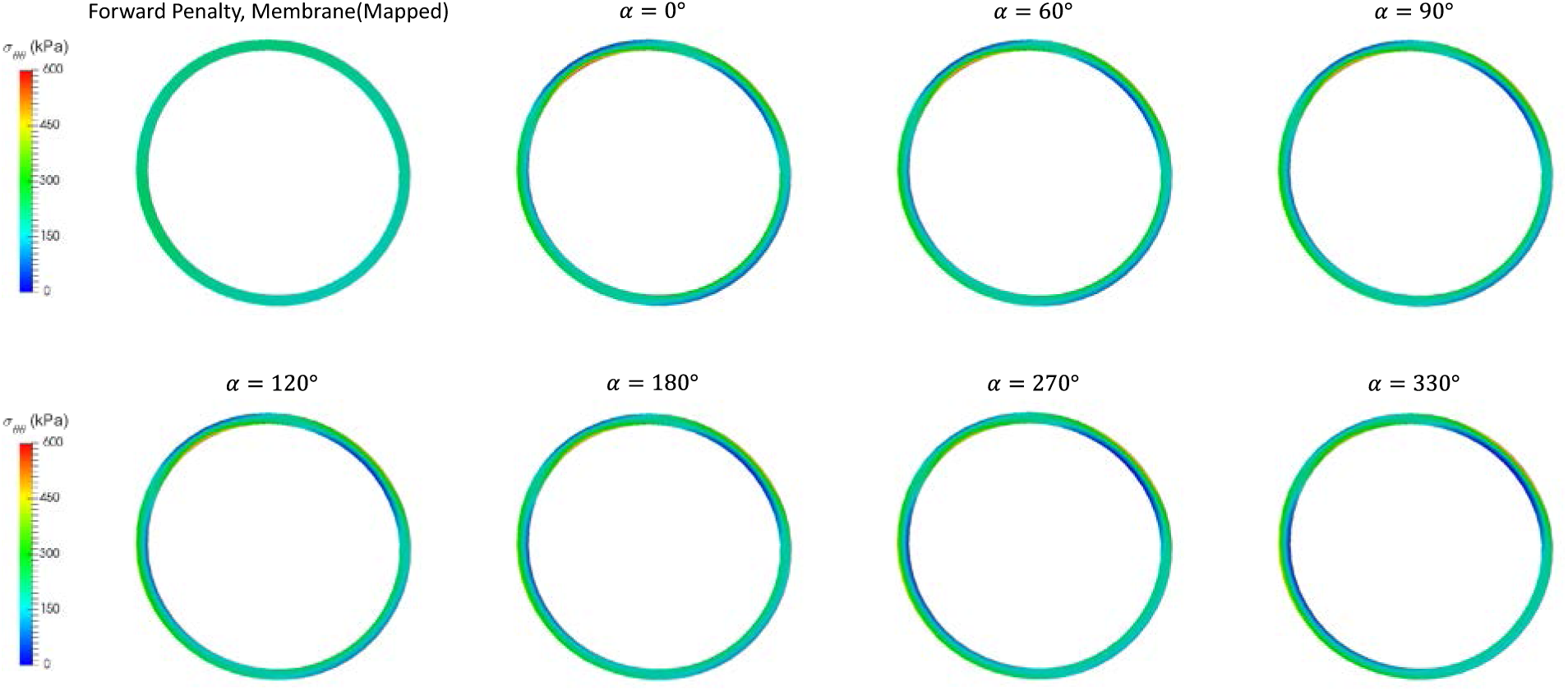
hoop stress in aortic rings using the forward penalty approach and the iterative approach (GPA) with different opening angles.

The probability density functions (PDFs) of the STPE are plotted in Figure 5. The PDFs are fitted using the Gaussian distribution. It can be observed that the PDF shifts leftward with the increase of the opening angle.

**Figure 5.**
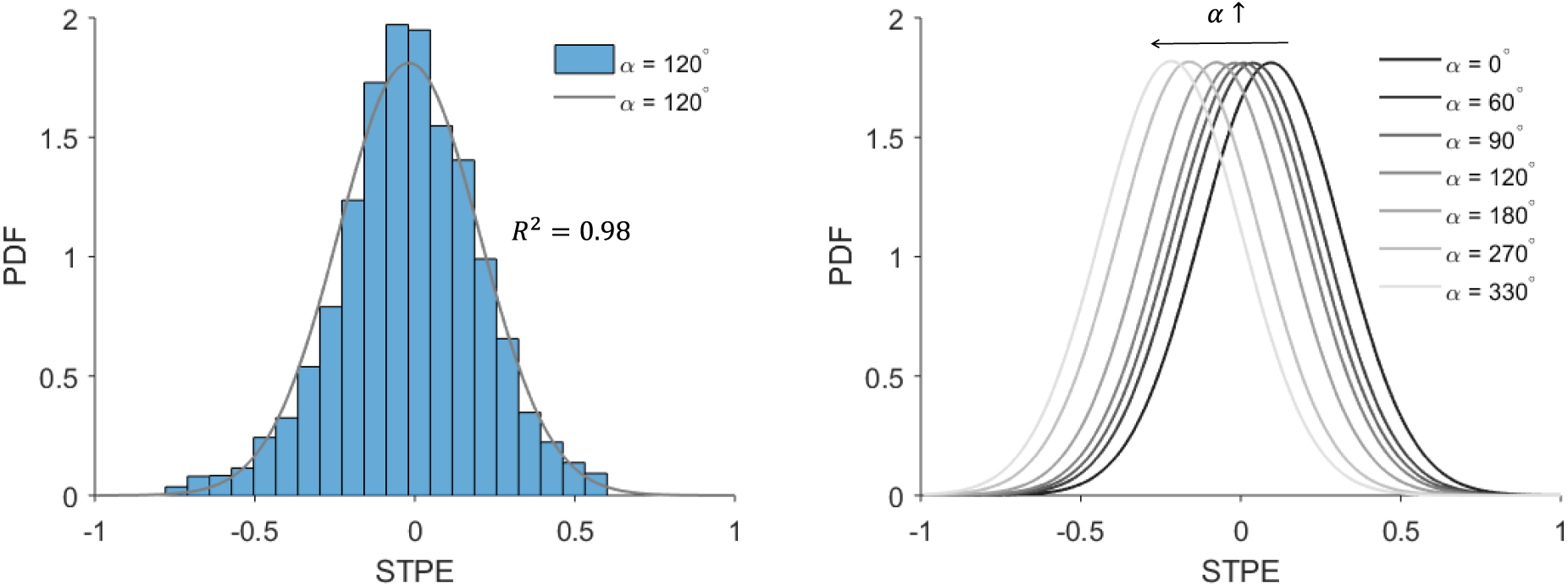
the probability density function (PDF) of the STPE is shown in the histogram and fitted using the Gaussian distribution (left) and fitted PDFs correspond to different opening angles (right).

#### 3.2.3 A Thick-walled Model with Layer-Specific Three-Dimensional Residual Deformation

In this section, the deformation gradient tensors 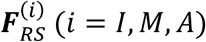 of Section 2.2 was incorporated in a FE simulation using the GPA. The ratio of intima, media and adventitia (18.53%, 45.56%, 35.91%) and the layer-specific GOH parameters (shown in Table 6) were taken from the median experimental value for human thoracic aortas in (Weisbecker et al. 2012). Layer-specific material parameter data for ATAA is also available in (Sassani et al. 2015; Sokolis et al. 2012). The geometrical parameters determining the residual deformation of abdominal aorta from (Holzapfel and Ogden 2010), same as Section 2.2 (Table 4), were directly used for the ATAA patient. (Sokolis 2015) documented layer-specific residual stretch and opening angle data for ATAA. Unfortunately, it is not compatible with the current three-dimensional residual stress model (Holzapfel and Ogden 2010). Specifically, (Holzapfel and Ogden 2010) considered different geometries of reference configurations for different layers and would need more complicated experimental setups.

**Table 6.** layer-specific GOH material parameters from (Weisbecker et al. 2012).

| | $C_{10}$ (kPa) | $k_1$(kPa) | $k_2$ | $\kappa$ | $\theta(^{\circ})$ |
| --- | --- | --- | --- | --- | --- |
| Intima | 17 | 4340 | 13.32 | 0.20 | 46.5 |
| Media | 14 | 140 | 11.90 | 0.21 | 38.4 |
| Adventitia | 10 | 390 | 6.79 | 0.23 | 52.3 |

Regardless of the discrepancy of the stress in the thickness direction (Figure 6, first row), the transmural mean stress field predicted by the forward penalty approach and the GPA are, again, almost identical, with a MAPE of 3.98% (Figure 6, second row). Since the details of transmural distribution of hoop stress is not clearly shown in the first row of Figure 6, we use Figure 7 to show hoop stress distributions in a ring predicted by method described in Section 3.1 (forward, membrane), Section 3.2.2 (opening angle *α =* 180°) and Section 3.2.3 (layer-specific 3D residual deformation), respectively.

**Figure 6.**
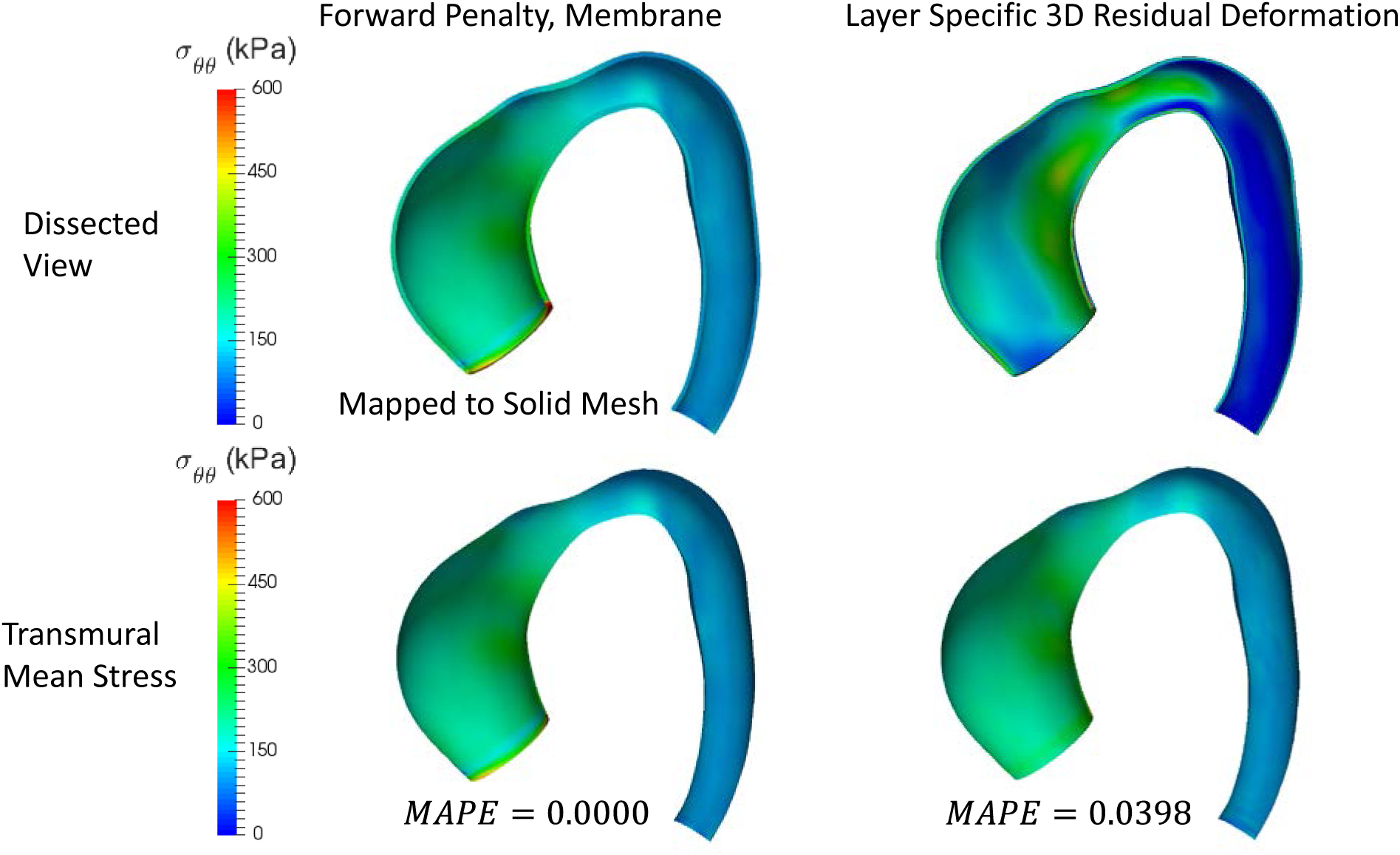
predicted results using the forward penalty approach and the iterative approach (GPA) with layer-specific three-dimensional residual deformations: (1) the hoop stress distribution in the dissected view (row 1), (2) the transmural mean hoop stress (row 2).

**Figure 7.**
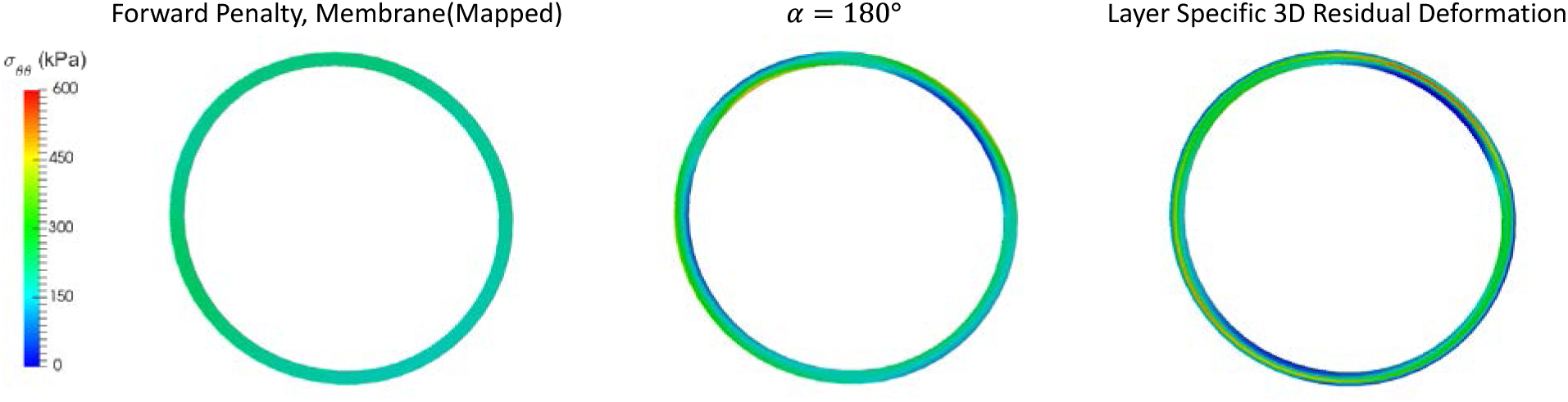
the hoop stress distribution in the aortic rings using the forward approach, the opening angle method (*α* = 180°) and the layer-specific 3D residual deformation.

## 4 Discussions and Conclusions

One of the biggest obstacles in the field of biomechanical analysis of the aorta is the difficulty in obtaining both the patient-specific material properties and the patient-specific residual deformations from *in vivo* clinical images. This paper offers an appealing prospect that the mean hoop stress (or hoop wall tension) of the aortic wall can be computed without knowing the mechanical properties and the residual deformations of the aortic tissue. Computation of patient-specific mean hoop stress can be greatly simplified by using membrane model and the forward penalty approach, which may be clinically valuable. In some wall strength tests (Ferrara et al. 2016; Pham et al. 2013), the intact wall is tested without separation of each individual layer, which corresponds to the averaged wall strength across the wall thickness, consistent with the membrane assumption. The mean hoop stress may be used together with the experimentally-obtained strength to calculate an approximation of rupture risk such as the rupture potential index (RPI) (Vande Geest et al. 2006).

Because of the difference in constituents and thus mechanical properties, the hoop stress distribution may not be uniform in multi-layer models. The iterative approaches such as the GPA, may yield detailed results with through-thickness and layer-specific stress distributions using multilayered thick-walled models. Therefore, it would be natural to combine layer-specific wall stress distribution with available layer-specific wall strength data (Sokolis et al. 2012) for a more detailed rupture/dissection analysis. Nonetheless, residual deformations are shown to be highly patient-specific and axial location-dependent (Sokolis 2015). Elastic properties also exhibit regional (Iliopoulos et al. 2009; Sassani et al. 2015) and intra-patient (Martin et al. 2015) variations. Thus, such complex patient- and layer-specific residual deformation and elastic property fields need to be noninvasively estimated for an accurate modeling prediction of clinical events (e.g. rupture). Currently, it is impossible to estimate the layer-specific and heterogeneous material and residual deformation parameters simultaneously from *in vivo* clinical images. We admit that hoop stress within each layer may be more useful than mean hoop stress for predicting some clinical adverse events such as aortic dissection. However, the mean hoop stress is clinically valuable too because it is patient-specific, which does not depend on material parameters and residual deformations.

The inclusion of residual deformation often reduces the hoop stress gradient, and thus tends to homogenize the hoop stress distribution in the *in vivo* deformed configuration (Chaudhry et al. 1997; Chuong and Fung 1986; Fung 1991; Holzapfel et al. 2000; Humphrey 2002; Raghavan et al. 2004). This makes the thin-walled hoop stress, or the mean hoop stress more physiologically relevant in the sense that it represents the ideal homogenized wall stress in single layer models. Homogenized stress state is an assumption for some growth models, e.g., (Polzer et al. 2013), and the method proposed in (Schröder and Brinkhues 2014) is based on smoothing the stress gradient. In this study, the incorporation of opening angles also tends to homogenize the hoop stress distribution as shown in Figure 1. In Figure 3, the MSTPE is close to 0 when 120~180 degree opening angle is incorporated. However, this value seems to be lower than the average value obtained from experiment (Sokolis 2015). This might be due to the assumption of uniform material properties and uniform thickness in the computational model, which could impact the transmural stress distribution. We also notice that a wide range of opening angle is documented in (Sokolis 2015), 120~180 degree opening angle can be considered feasible.

The transmural mean axial/longitudinal stress of the aorta may be statically determinant when the longitudinal force is known. The ascending aorta also has *in vivo* longitudinal deformations/stretches due to the heart movements during cardiac cycles. Such boundary condition is very complex and it can be difficult to model in a FE simulation. In the present study, a simplified boundary condition was used: the boundary nodes were only allowed to move in the radial directions. We have tried different boundary conditions such as prescribing the longitudinal forces, but encountered convergence problems in the FE simulations. The *in vivo* longitudinal boundary conditions would significantly impact the longitudinal stress field, which warrants further studies in the future.

In conclusion, due to static determinacy, the transmural mean hoop stress in the *in vivo* configuration of the aorta is independent of mechanical properties and residual deformations. The forward penalty method, which enforces a rigid condition as the penalty treatment, can greatly simplify the computation of the mean hoop stress for patient-specific geometries.

## 5 Acknowledgements

This study is supported in part by NIH grants HL104080 and HL127570. Liang is supported by an American Heart Association postdoctoral fellowship 16POST30210003.

## 6 Conflict of Interest

The authors declare that they have no conflict of interest.

